# Ultrasound-actuated drug delivery with acoustic percolation switches

**DOI:** 10.1101/2024.05.10.593654

**Authors:** Maria Paulene Abundo, Anna T. Tifrea, Marjorie T. Buss, Pierina Barturen-Larrea, Zhiyang Jin, Dina Malounda, Mikhail G. Shapiro

## Abstract

Devices that can be remote-controlled under image guidance to precisely deliver biomedicines to sites of disease are a major goal of biomedical research. However, most existing externally triggered delivery systems are based on complex micromachines that are controlled with electromagnetic waves and require custom external instrumentation. Here we present a drug delivery platform comprising a simple protein-containing hydrogel that can be both imaged and triggered to release drugs at specific locations using widely available ultrasound imaging devices. This technology is based on the addition of air-filled protein nanostructures called gas vesicles (GVs) to hydrogel delivery vehicles. While intact, GVs sterically block the release of drug payloads and allow the vehicle to be imaged with ultrasound. An increase in ultrasound pressure causes the collapse of GVs within hydrogels present at the desired anatomical location, instantly creating percolation channels and triggering rapid drug release. Both the imaging and release are performed using a common diagnostic ultrasound probe. We implement this concept by establishing ultrasound-controlled drug diffusion and release from hydrogels *in vitro* and demonstrating targeted image-guided protein delivery *in vivo* following oral administration. We use this approach to deliver anti-inflammatory antibodies to treat gastrointestinal inflammation in a rat model of colitis. Targeted acoustic percolation switches (TAPS) open a conduit for local, image-guided drug delivery with a simple formulation and commonplace ultrasound equipment.

---

The development of technologies for remote-controlled, targeted drug delivery represents a ‘holy grail’ of biomedical research. When implemented successfully, these platforms allow therapeutics to reach specific anatomical locations in complex tissues such as the gastrointestinal (GI) tract, providing efficacy at the intended site of action while minimizing side-effects. Ideally, remote-controlled drug delivery would be performed under image guidance, making it possible to monitor vehicle location, trigger release, and confirm payload delivery. However, executing these actions *in vivo* remains a challenge. Most existing approaches utilize complex micromachined devices triggered by electric^1^ or magnetic fields^2,3^, which are challenging to focus *in vivo*; or by light^4,5^, which has limited tissue penetration^6^. Moreover, these devices tend to be non-degradable^7^, expensive to produce, difficult to miniaturize and require dedicated external apparatus for monitoring and control^8,9^.

If instead it were possible to develop a drug delivery vehicle that could be tracked and triggered with diagnostic ultrasound, this would enable image-guided drug delivery using ubiquitous, low-cost instruments with deep penetration (> 10 cm) and high resolution (< 1 mm) ^10^. Moreover, if such vehicles could be produced via the simple mixing of biocompatible components, this would accelerate their development for a broader range of clinical applications.

Here we introduce a new class of hydrogel materials that respond to diagnostic ultrasound to enable targeted, image-guided drug delivery. To interact with sound waves, these materials use a unique class of air-filled protein nanostructures called gas vesicles (GVs) derived from buoyant photosynthetic microbes^11,12^. GVs comprise a 2.4 nm-thick protein shell that encloses a hollow, air-filled compartment with dimensions on the order of 100 nm^11,13–15^ (**Fig. 1, a-b**). Due to their ability to scatter sound waves, GVs have been developed as contrast agents for ultrasound imaging^16–21^. However, they have not been used in drug delivery devices.

**Figure 1.**
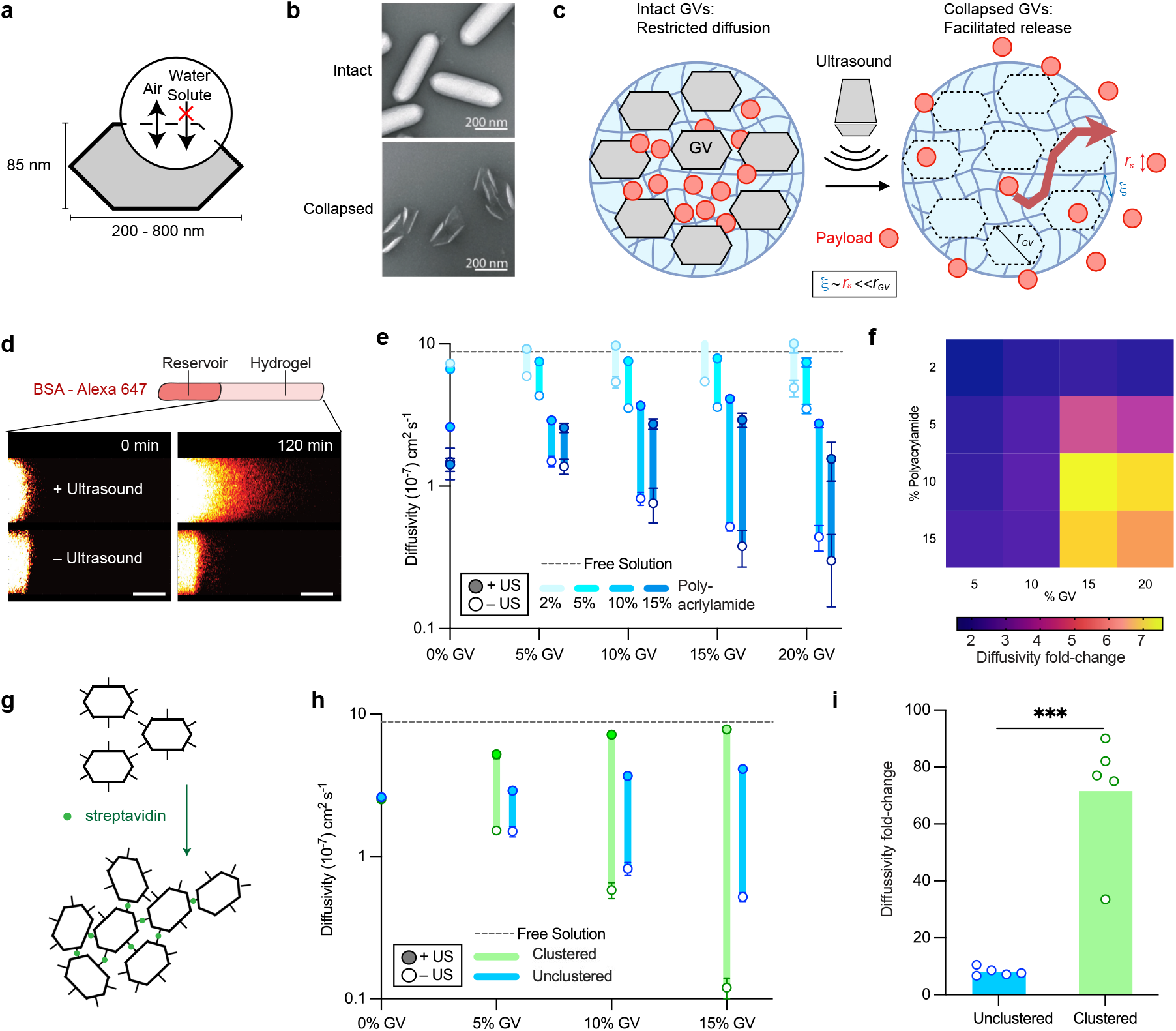
Gas vesicle-containing hydrogels produce ultrasound-modulated diffusivity changes. (**a**) Schematic of a gas vesicle highlighting its impermeability to surrounding molecules. (**b**) TEM images showing intact (left) and pressure-collapsed (right) gas vesicles. (**c**) Diagram of the TAPS concept. Intact GVs block payload diffusion. Collapse of the GVs with a diagnostic ultrasound pulse turns them into percolation channels, allowing rapid payload release. (**d**) Illustration of the experimental set-up for measuring the diffusion of BSA-AlexaFluor 647 in GV-containing hydrogels. (**e**) Diffusivity of BSA-AlexaFluor through GV-gels of varying GV and polyacrylamide volume fractions with and without ultrasound exposure (*N* = 5, *mean* ± *s*.*e*.*m*.). (**f**) Fold-change in diffusion between the intact GV ‘ultrasound off’ and collapsed GV ‘ultrasound on’ states for different gel compositions. (**g**) Schematic of GV clustering using biotin and streptavidin. (**h**) Diffusivity changes of BSA-AlexaFluor through gels containing clustered and unclustered GVs with and without ultrasound exposure for 10% polyacrylamide and varying GV volume fractions (*N* = 5, *mean* ± *s*.*e*.*m*.). (**i**) Fold-change in diffusivity upon ultrasound exposure in 10% polyacrylamide gels containing a 15% volume fraction of clustered and unclustered GVs (*p* = 0.0002, *N* = 5).

We hypothesized that a drug delivery approach could be implemented by taking advantage of the ability of GVs to collapse irreversibly under well-defined acoustic pressure (above a sharp threshold typically ∼ 500 kPa^17^), causing them to lose more than 85% of their volume^13,22^ (**Supplementary Table 1**). We designed a hydrogel vehicle incorporating GVs alongside a protein payload, such that the presence of the drug-impermeable intact GVs sterically restricts drug diffusion, while ultrasound-triggered GV collapse instantly converts the GVs into percolation channels, leading to rapid drug release (**Fig. 1c**). Before triggering this release, the hydrogels can be imaged with lower-amplitude ultrasound to visualize their location inside the body, while GV collapse leads to a loss of ultrasound contrast^16,23,24^ providing confirmation that release has been triggered.

In this study, we implemented this concept by generating hydrogels containing GVs and protein payloads. We performed detailed measurements of ultrasound-triggered diffusion and release, using the resulting insights to construct targeted acoustic percolation switches (TAPS). We demonstrated the ability of these devices to release proteins in the lower gastrointestinal tract following oral administration and ultrasound actuation and used them to deliver an anti-inflammatory antibody drug to the duodenum in a rat model of colitis, resulting in reduced inflammation and improved health measures. TAPS enable local, image-guided and imaging-confirmed drug delivery using a simple formulation and ubiquitously available ultrasound imaging equipment.

## GV-containing hydrogels exhibit ultrasound controlled biomolecular diffusivity

We tested the basic hypothesis that the presence of GVs in a hydrogel would restrict the diffusion of other biomolecules by preparing polyacrylamide gels with GVs and measuring the diffusivity of fluorescently-labeled bovine serum albumin (BSA). We focused on biomolecular payloads because they represent potent therapeutics that are challenging to deliver to locations such as the GI tract. We polymerized gels containing different volume fractions of GVs (purified from *Anabaena flos-aquae*) and acrylamide/bis-acrylamide inside glass capillaries and loaded an adjacent reservoir with a solution of BSA-AlexaFluor 647. Then we used confocal microscopy to track the movement of fluorescent signal through the capillary ^25^ (**Fig. 1d**) and fitted the resulting curves to a one-dimensional diffusion equation (**Supplementary Fig. 1**).

We used a diagnostic ultrasound transducer operating at 6 MHz to collapse the GVs in a subset of the gels. Our results confirmed that intact GVs restrict protein diffusion relative to GV-free hydrogels (**Fig. 1e, empty circles**), and that acoustic collapse of the GVs leads to a large increase in this diffusivity (**Fig. 1e, filled circles**). The dynamic range of diffusivity between the ultrasound ‘on’ and ‘off’ conditions was optimal at GV and polymer concentrations of 15% and 10% by volume, respectively (**Fig. 1f**). This is consistent with the need for low drug diffusion through the background gel, high steric blockage by intact GVs and the ability of collapsed GVs to create interconnected percolation channels.

Building on these results, we hypothesized that the dynamic range of our hydrogels could be widened by clustering the GVs prior to gelation. The resulting increase in inter-GV connectivity would make diffusion around intact GVs more challenging while enhancing the connectivity of the post-collapse percolation network. To test this hypothesis, we clustered GVs using biotin and streptavidin^16,26^, generating aggregates with hydrodynamic diameters of around 1 μm (**Fig. 1g**). Embedding these pre-clustered GVs in the hydrogel at volume fractions of 10-15% substantially reduced BSA diffusivity compared to unclustered GVs, while boosting the diffusivity upon GV collapse (**Fig. 1h**). The optimized composition of 15% clustered GVs and 10% acrylamide exhibited a pressure-induced diffusivity fold-change of 71.5 ± 9.8 (mean ± SEM, **Fig. 1i**).

## TAPS enable imaging and spatially targeted drug release *in vitro*

Having demonstrated switchable diffusivity, we set out to test the ability of GV-containing gels to release drug payloads on command. We formed prototype TAPS devices by mixing hydrogel-forming reagents, clustered GV and fluorescent BSA in 3D-printed cylindrical molds with a diameter of 0.5 mm and a height of 2 mm (**Fig. 2a**). The gels were then preincubated in a phosphate-buffered saline (PBS) solution for 2 hours to remove loosely bound BSA on or near the hydrogel surface and transferred into a stirred vial containing fresh media (**Fig. 2b**). We applied ultrasound to a subset of the gels and tracked the fluorescence intensity of the solution over 12 hours, finding that while TAPS gels retained most of their payload in the absence of ultrasound treatment, devices actuated by ultrasound released most of their BSA within 2 hours and reached nearly complete release in 5 hours (**Fig. 2c**).

**Figure 2.**
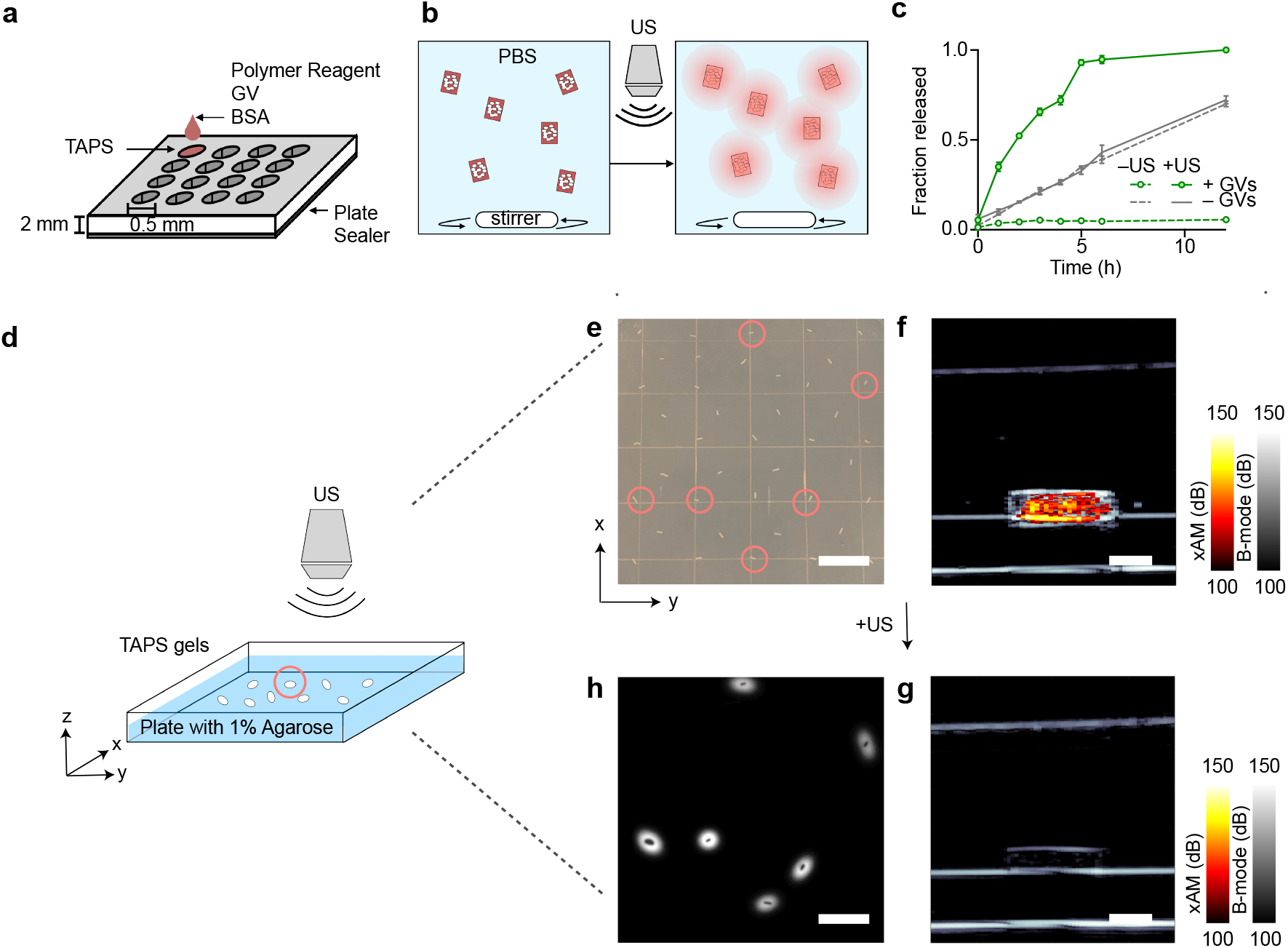
TAPS enable spatially selective ultrasound-guided, ultrasound-actuated and ultrasound-confirmed payload release. **a**, Schematic of the 3D-printed mould used to form millimetre-sized TAPS cylinders. **b**, Diagram of experimental set-up used to track fluorescent payload release over time. GV-gels are placed into a vial containing phosphate buffered saline (PBS) and stirred for 12 hours. The kinetics of payload release is monitored by aliquoting samples from the vial every hour and determining the fluorescence intensity. **c**, Release profiles of clustered TAPS (green) with and without ultrasound exposure *(N* = 3, *mean* ± *s*.*e*.*m*.) compared to the no-GV hydrogel only controls (grey). Fraction released is defined as the fluorescence signal in the vessel normalized to the fluorescence signal at the end of the experiment. **d**, Diagram of the experimental setup used to assess spatial targeting performance of the drug delivery system *in vitro*. TAPS are dispersed on a 1 %wt agarose phantom, and an ultrasound probe is used to target individual gels for collapse. **e**, Top-down photo of the plate showing dispersed TAPS in agarose. Devices circled in red depict those selected for ultrasound targeting at t = 0. **f**, xAM ultrasound image of a representative TAPS prior to ultrasound actuation. **g**, xAM image of the same TAPS after ultrasound activation. **h**, Fluorescence subtraction image obtained between t = 12 hours post GV collapse and at t = 0. Scale bars: **e**,**h**,10 mm; **f**,**g**, 1 mm.

Having demonstrated payload release from these model TAPS, we tested their ability to be imaged with ultrasound and actuated in a spatially selective manner. We dispersed our devices in 1% agarose with inter-device spacing of approximately 5 mm (**Fig. 2d**). We used a diagnostic ultrasound transducer operating at 6 MHz and 200 kPa peak positive pressure to locate and image the gels using a GV-selective nonlinear xAM pulse sequence^27^ (**Fig. 2f**). The gels were readily visible under ultrasound. We then selected several individual TAPS gels (indicated by the dashed red circles in **Fig. 2e**) for actuation by increasing the ultrasound transmit pressure to > 550 kPa, leading to GV collapse, as confirmed immediately by the disappearance of their xAM ultrasound contrast (**Fig. 2g**). We observed that this remote actuation of the gels resulted in selective payload release, as shown by the localized diffusion of fluorescent BSA into surrounding agarose (**Fig. 2h**). These results validate the fundamental capability of TAPS to be used as ultrasound-imaged, ultrasound-actuated and ultrasound-confirmed delivery vehicles.

## TAPS enable spatially selective ultrasound-triggered drug release *in vivo*

One of the most immediate applications of spatially triggered drug delivery devices is the gastrointestinal tract, due to a range of GI pathologies requiring local treatment^28^. Thus, we chose the GI tract as the initial proving ground for TAPS. To demonstrate the spatiotemporal control of drug release in vivo, we used gavage to orally administer a suspension of BSA-loaded TAPS in PBS to Sprague Dawley rats (**Fig. 3a**). After 5 hours, we used a diagnostic ultrasound probe to locate GV-gels in the ileum and cecum (**Fig. 3, b-c** and **Supplementary Fig. 2**), using GV-selective BURST imaging for maximum detection sensitivity^23^. We then triggered payload release by increasing the transmit pressure to > 550 kPa and confirmed GV collapse by the disappearance of BURST signal (**Fig. 3d**). We then collected the large intestines of the rats and laid them out in PBS. Following established protocols^29,30^, we gently flushed the intestines with PBS after 12 hours to evacuate TAPS and any solid debris, leaving behind fluorescently labeled protein that was released from the gels and absorbed by intestinal walls. Fluorescence images of the GI tract showed significant signal localized in the proximal colon only in rats that received the release-activating ultrasound pulse (**Fig. 3, e-f**). These results show that TAPS can be imaged and triggered noninvasively with ultrasound inside intact living animals, leading to localized release of a protein payload.

**Figure 3.**
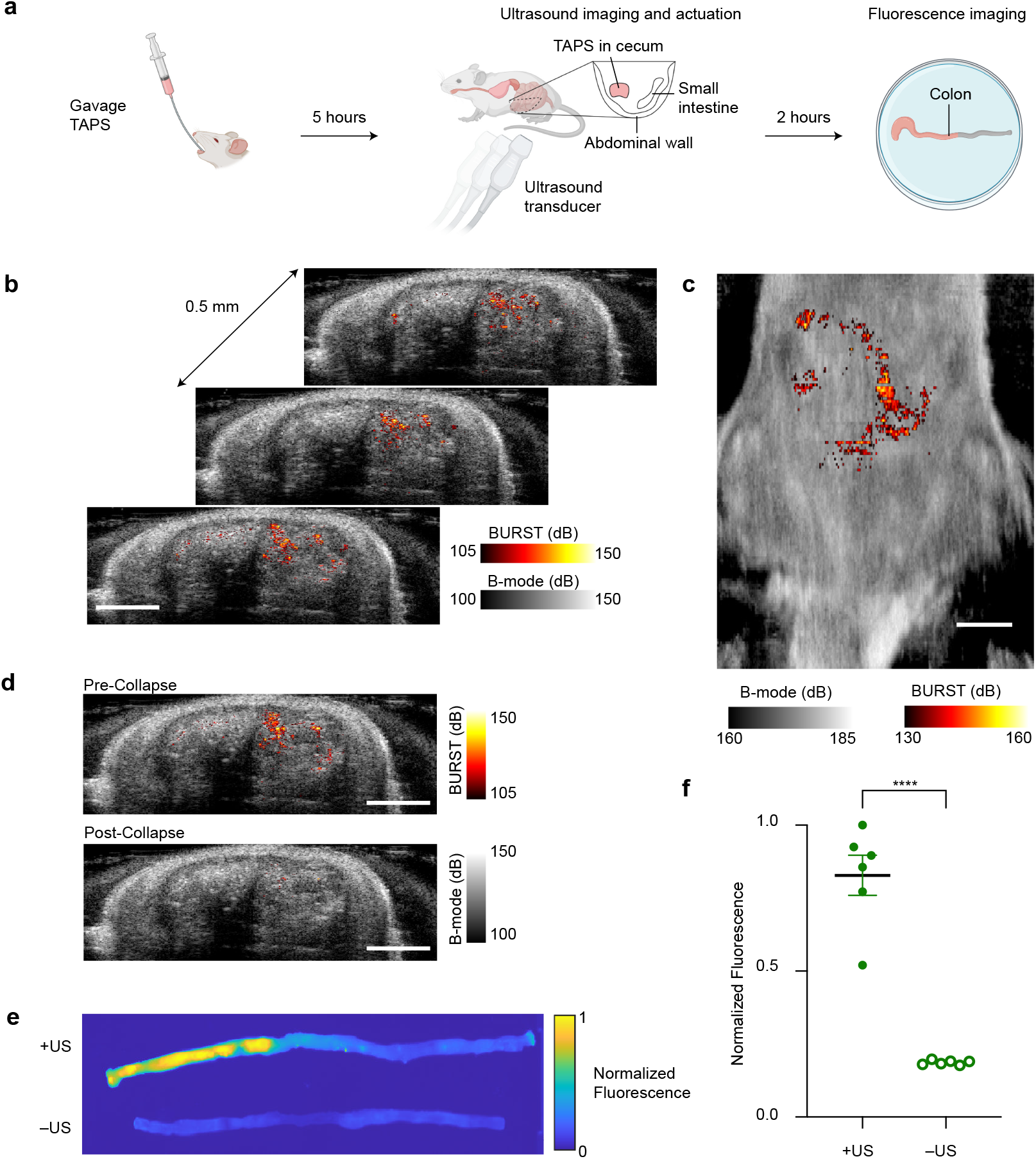
Ultrasound imaging and targeted payload release from TAPS in the gastrointestinal tract. **a**, Illustration of the in vivo imaging and release experiment. **b**, Representative stack of coronal BURST ultrasound images (heat color map) of the lower abdomen area of a rat overlaid on anatomical B-mode images (grayscale) five hours after TAPS gavage. BURST signal was not observed in rats that were given saline (**Supplementary Fig. 2**). **c**, Representative top-down projection of BURST and B-mode signal across all the imaged planes. **d**, Representative ultrasound images of the cecum pre-actuation (top) and post-actuation of release from the TAPS gels. **e**, Representative fluorescence images of colons from rats gavaged with TAPS, with and without ultrasound activation of payload release. **f**, Retained fluorescence signal in the colons of TAPS-gavaged rats with and without ultrasound actuation. *P < 0*.*0001*. Scale bars: **b**-**d**, 1 cm.

## TAPS release of etanercept in the GI tract enables the treatment of colitis

Having established the basic functionality of TAPS *in vitro* and *in vivo*, we set out to demonstrate their application to delivering a protein therapy in a preclinical disease model. Inflammatory bowel diseases, including ulcerative colitis, affect more than 2 million people in the United States^31^. Patients with colitis often require lifelong treatment and pain management, experiencing reduced quality of life due to episodes of relapse characterized by moderate to severe lesions in the large intestine. Among other treatments, colitis is treated with biologics inhibiting the pro-inflammatory cytokine tumor necrosis factor (TNF)^32^. These drugs are administered systemically, typically requiring high systemic doses to reach therapeutic concentrations at the sites of GI inflammation. The resulting systemic immunosuppression can lead to side-effects such as infections and tumorigenesis^33–35^. If anti-TNF therapies could be delivered more directly to the colon, this would increase their dose at the target site while mitigating systemic side-effects. We hypothesized that TAPS would allow us to deliver the monoclonal anti-TNFα antibody etanercept to the colon and thereby treat colitis in a rat model of the disease. Etanercept is FDA-approved for the treatment of autoimmune diseases and has previously been validated to treat experimental colitis in rodents due to its cross-reactivity with rat TNFα^36–38^.

To formulate TAPS with etanercept, we incorporated the drug and GVs in calcium alginate – a hydrogel validated for sustained release of monoclonal antibodies^39^. To protect the hydrogels from stomach pH and enzymes, we dip-coated them with Eudragit FL30D55 (**Fig. 4a**). The ultrasound-mediated protein release performance of this system was validated by releasing fluorescent BSA in simulated rat gastric fluid (**Fig. 4b**).

**Figure 4.**
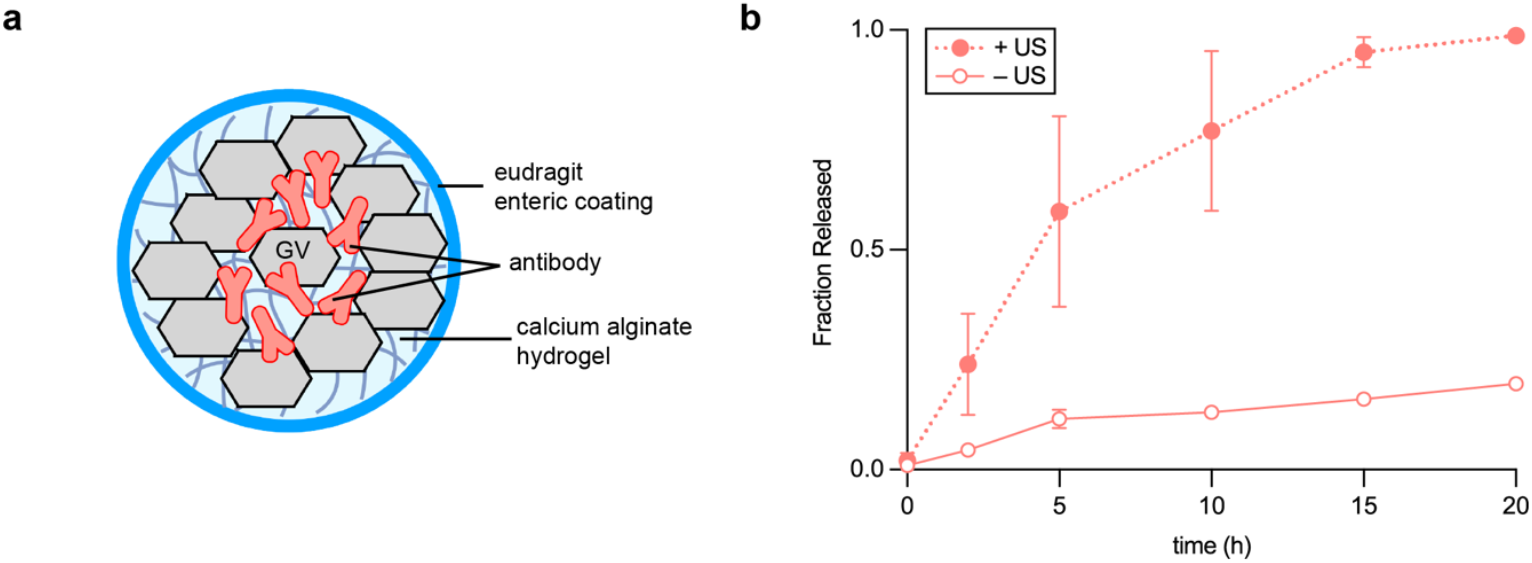
TAPS design for oral delivery of biomolecules. **a**, Illustration of antibody-loaded TAPS based on calcium alginate. TAPS for oral gavage in rodents are coated with a thin enteric coating layer of Eudragit. **b**, Ultrasound-mediated controlled release kinetics of BSA-AlexaFluor 647 from calcium alginate-based TAPS.

We induced severe colitis in Sprague-Dawley rats by adding 4 percent dextran sulfate sodium (DSS) to their drinking water for five days (**Fig. 5a**), confirming that DSS-treated rats lost weight relative to untreated controls (**Supplementary Fig. 3**). We treated the animals with daily oral gavage of TAPS-etanercept at a dose of 12.5 mg (selected based on a preliminary dose-ranging experiment, **Supplementary Fig. 4**) from days 6 to 11. In a subset of subjects, at 5 hours after gavage, when the gels are expected to be in the cecum (**Fig. 4, b-c**), we triggered etanercept release using a diagnostic ultrasound transducer scanned over the abdomen with transmit pressure > 550 kPa (**Fig. 5b**). Control conditions included rats that were administered TAPS-etanercept but not ultrasound and rats that were treated with ultrasound but no TAPS. Before and during the treatment window we measured the rats’ weight and disease activity index (DAI)^40^, an aggregate measure encompassing stool consistency, fecal bleeding and weight loss. The subjects were euthanized at day 12 for blinded histopathological analysis of the colon by a nonaffiliated pathologist.

**Figure 5.**
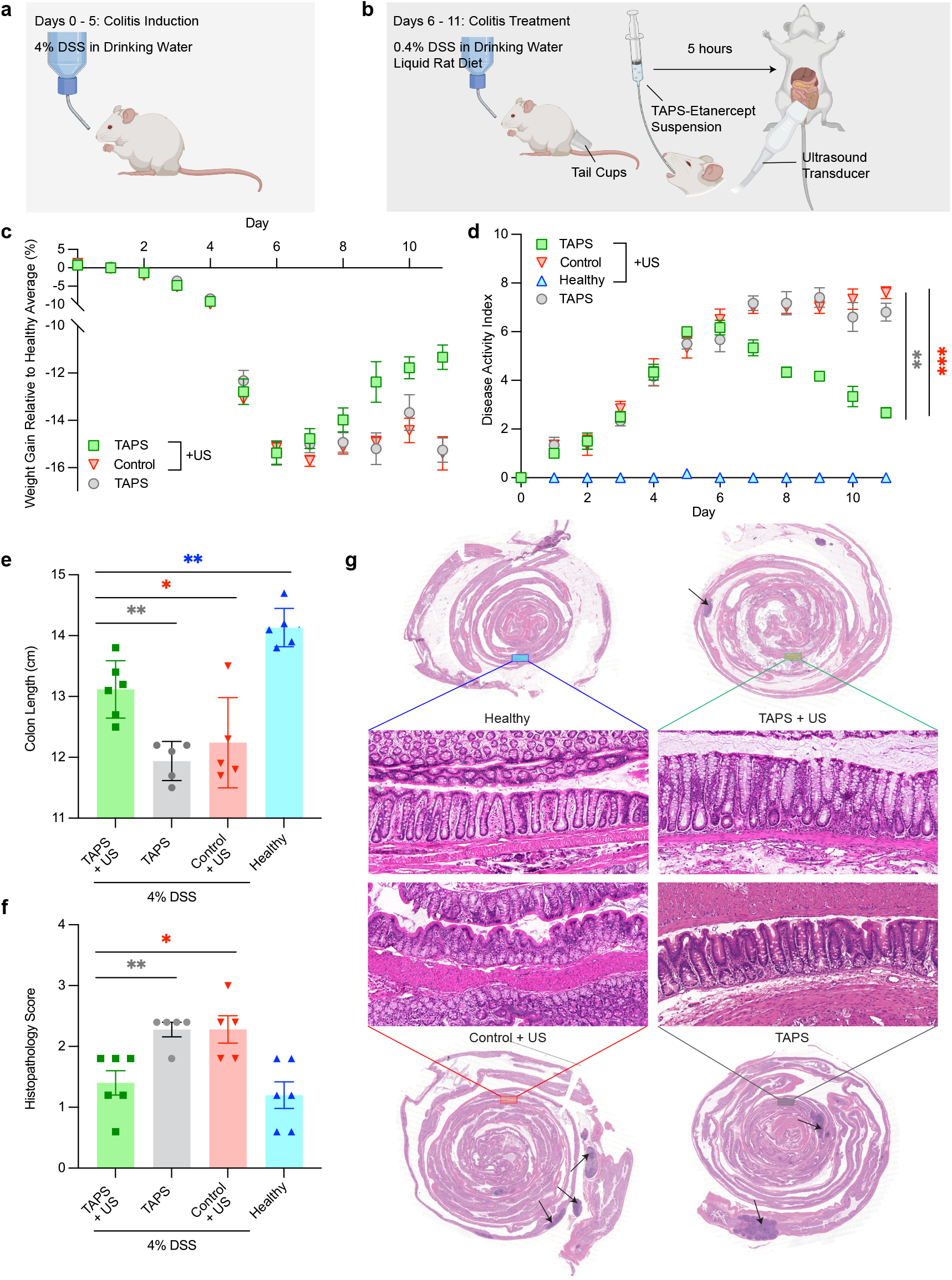
TAPS with etanercept enable ultrasound-triggered treatment of colitis in a rat model. **a-b**, Experimental workflow used to demonstrate treatment of colitis in rats using the TAPS delivery vehicles. **c**, Change in rat weight gain rate relative to healthy controls over the treatment period for rats with severe colitis that were given TAPS treatment or controls. Where not seen, error bars are smaller than the symbol. The p-values for the statistically significant points during the treatment period are as follows for (i) TAPS+US vs TAPS: Day 9, *P = 0*.*0343*; Day 11, *P = 0*.*0004* (ii) TAPS+US vs control: Day 9, *P = 0*.*01783*; Day 10, *P = 0*.*0035*; Day 11, *P = 0*.*0010*. **d**, Time course of the Disease Activity Index (DAI), which is the sum of the stool consistency index (0–3), fecal bleeding index (0–3) and weight loss index (0–4). The p-values for the statistically significant points during the treatment period are as follows for (i) TAPS+US vs TAPS : Day 7, *P = 0*.*0023*; Day 8, *P = 0*.*0003*; Day 9, *P = 0*.*00002* ; Day 10, *P = 0*.*0013*; Day 11, *P < 0*.*00001* (ii) TAPS+US vs control: Day 7, *P = 0*.*0027*; Day 8, *P = 0*.*00001*; Day 9, *P < 0*.*00001*; Day 10, *P = 0*.*000053*; Day 11, *P < 0*.*00001*. **e**, Colon length of rats after the indicated treatments. The p-values are (vs TAPS+US) (i) TAPS: *P = 0*.*0011* (ii) control: *P = 0*.*0405* (iii) healthy: *P = 0*.*0013* **f**, Histopathology scores for rats in (b) and a no-DSS health control group. The p-values are (vs TAPS+US) (i) TAPS: *P = 0*.*0060* (ii) control: *P = 0*.*0166*. **g**, Representative H&E-stains of the colonic sections of different rat groups after the indicated treatment. Black arrows are pathologist-identified abscessed regions.

In rats that received ultrasound actuated TAPS-etanercept treatment, we observed a significant recovery of body weight (**Fig. 5c**), reversal of the DAI (**Fig. 5d**), recovery of colon length (**Fig. 5e**) and reduced colon tissue damage (**Fig. 5, f-g**) compared to controls. Post-mortem histological imaging revealed that the treated group had mild to no histological alterations by the end of the treatment, with regular structure of the lamina propria restored and comparable to those in the healthy rat controls. In contrast, rats that were administered TAPS without ultrasound actuation or ultrasound alone had crypt abnormalities and inflammatory acute infiltration of the sub-epithelium and lamina propria (**Fig. 5g**). Together, these results demonstrate the capability of TAPS to deliver an efficacious biologic disease treatment to the GI tract following oral administration.

## DISCUSSION

This work establishes TAPS as a unique drug delivery system allowing everyday diagnostic ultrasound devices to image, trigger and confirm the release of therapeutics at specific locations inside the body. TAPS leverages the ability of GVs to simultaneously act as ultrasound contrast agents and collapsible steric blockers, allowing simple hydrogels to be both visualized and remotely actuated with ultrasound.

In this study, we established the fundamental concept of a hydrogel in which drug diffusion is activated by ultrasound, demonstrated the ability of this concept to support remote-controlled release of biomolecular payloads, and provided a proof of concept for its use to deliver a biologic therapy to the GI tract. In future studies, the TAPS technology can be extended to a wider range of cargoes^41^ — from small-molecule drugs (∼ 1 nm) to viruses such as adeno-associated virus (>20 nm) — by tuning the hydrogel mesh size relative to the hydrodynamic radius of the payload. These payloads could be released in contexts ranging from ingestible pills to implantable and interventional medical devices or smart robots navigating through a tissue^42^. In addition, a combination of GV types with varying collapse pressures^17^ could be used to sequentially release multiple payloads. Furthermore, acoustophoresis could be used to pattern GVs within the material^43^ to further optimize release properties.

As with any new technology, further work is needed to optimize the TAPS system and enable its broadest range of clinical applications. Depending on the therapeutic payload, it may be necessary to use hydrogels other than the polyacrylamide and alginate used in this study and establish their compatibility with GVs. Additionally, for repeated administration of GV-containing hydrogels, it will be important to characterize the immune accessibility and potential immunogenicity of the GV proteins. Furthermore, for patients to be able to use TAPS at home, consideration must be given to ultrasound devices, including low-cost portable devices^44^, adhesive ultrasound^45–48^ and machine learning algorithms to automatically recognize and trigger delivery vehicles at specific anatomical sites. For uses in the GI tract, it may also be necessary to deal with confounding gas and solid contents in ultrasound images. This task will be aided by GV-specific ultrasound pulse sequences^23,27,49,50^ and patient preparation through diet.

Notwithstanding the need for further research, the relative simplicity and versatility of the TAPS approach and its compatibility with existing FDA-approved hydrogel materials, drugs and ultrasound devices will allow this technology to percolate toward valuable clinical applications.

## ACKNOWLEDGEMENTS

The authors thank Yuxing Yao and John Brady for helpful discussions. MPA was funded by the A*STAR graduate fellowship. MTB was funded by the National Science Foundation graduate research fellowship. This research was supported by the Jacobs Institute for Molecular Engineering in Medicine, the David and Lucille Packard Foundation and the Dreyfus Teacher-Scholar Award. MGS is an Investigator of the Howard Hughes Medical Institute.

## AUTHOR CONTRIBUTIONS

MPA and MGS conceived and designed the study. MPA, ATT, MTB, PBL and ZJ designed, planned and executed experiments. MPA, MTB and ATT analyzed the data. DM prepared gas vesicles. MPA, ATT and MGS wrote the manuscript with input from all authors. MGS supervised the research. All authors have given approval to the final version of the manuscript.

## MATERIAL AND METHODS

### Animals

5-week-old male Sprague Dawley rats weighing between 150-160g were purchased from Charles River (USA). The rats were housed in the animal facility of the Office of Laboratory Animal Resources (OLAR) at the California Institute of Technology with a 13h/11h light/dark cycle with cage conditions kept between 22 to 24°C and 30 to 70% humidity. The rats were monitored daily for overall well-being and given ad libitum access to water and standard laboratory chow. All animals were acclimated for three days prior to performing experiments that are in accordance with an approved Caltech Institutional Animal Care and Use Committee (IACUC) protocol.

### Preparation of gas vesicles

Purified GVs from *Anabaena flos-aquae* with GvpC removed (AnaΔC) and subsequently, clustered to form larger aggregates were prepared as previously described^16,22^.

### Diffusion characterization

50 μl of hydrogel reaction mixture is made by copolymerizing a predetermined volume fraction of acrylamide and bis-acrylamide (A7168: Sigma Aldrich) in prefiltered pH 7.4 phosphate buffered saline (PBS) solution at a fixed cross-linking density of 2.7%. 50 μl of purified *Anabaena Flos-Aquae* GVs solution was concentrated to a predetermined volume fraction (optical density, OD 24 = 1% vol. GV) and added to the hydrogel reaction mixture. The hydrogel reaction mixture is then degassed in a vacuum chamber for 30 min to rid the system of oxygen. 0.1 μl and 0.5 μl of initiators tetramethyl ethylene diamine (T9281: Sigma Aldrich) and 10% ammonium persulfate solution (A3678: Sigma Aldrich) respectively was then added to the reaction mixture and agitated slightly using a pipette to start the free-radical polymerization at room temperature (25°C). Using the prepared syringe-capillary system, the hydrogel reaction mixture is immediately withdrawn into the glass capillary, leaving about a 1 inch of the glass capillary empty at the top for dye loading later. This recipe is expected to produce hydrogels of approximately 19 nm^51^ pore sizes. The bottom of the glass capillary is quickly capped with some clay to prevent the hydrogel from drying or leaking out. To collapse the GVs in the GV-gel, the filled glass capillary is first placed into a PBS-filled container and held in place by solidified 1% Agarose. An L10-4V 128-element linear array ultrasound transducer (Verasonics) operating at peak positive pressure of 600 kPa was then held in place over the capillary for 3 mins to collapse the GVs in the gel. GV collapse is verified visually from the loss of gel opacity, and from the loss of ultrasound signal within the gel. The capillary loaded gels are then equilibrated at room temperature with PBS for at least 6 hours prior to experiments. Albumin from Bovine Serum (BSA) with AlexaFluor™ 647 conjugate (A34785: ThermoFisher Scientific) was diluted to 500 μg/ml using PBS and equilibrated at room temperature for 5 mins. A 1 ml BD Luer-Lok™ tip syringe fitted with a blunt 30G needle (SAI Infusion Technologies) was then used to inject the protein dye near the interface of the hydrogel casted within a 3-inch open-ended glass capillary (1B120-3: World Precision Instruments), ensuring all the air bubbles are extracted prior to experiments. Both ends of the capillary are then capped with clay to prevent evaporation. The experiments were then carried out as described^25^ for rapid measurements of protein diffusion through hydrogels using a Zeiss LSM 800 inverted microscope fitted to a 2.5x objective lens (numerical aperture: 0.075). Fluorescence imaging was performed under an Alexa 647 channel (excitation: 587/25 nm, emission: 647/70 nm) and images were taken every 6 mins over a period of 2 hours.

The time series of fluorescence images from the capillary diffusivity experiments capillary were imported into MATLAB for image processing. The time evolution of the fluorescence intensity across the capillary length from the fluid-gel interface was recorded and averaged across the capillary cross-sectional area. Since the length of the gel (5 cm) is significantly larger that its diameter (0.6 mm) one can approximate the protein-dye transport in the gel to follow a 1-D diffusion model through an infinite slab given by the following equation:

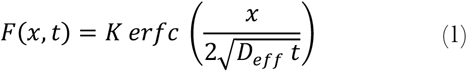

where *K* and *D*_*eff*_ are constants that represent the gel partitioning and diffusion coefficients of BSA respectively, *F* is the fluorescence intensity normalized to the dye reservoir intensity, *x* is the distance from the interface and *t* is the time. The experimental data was then fitted to equation (1) using a nonlinear programming solver on MATLAB to obtain the two constants.

### Acrylamide TAPS preparation

The acrylamide TAPS cargoes utilized the same hydrogel recipe as described in the BSA diffusivity experiments but with the addition of BSA-AlexaFluor 647 at a final dye concentration of 500 μg/ml also mixed into the polymerization precursor media. The reaction mixture was then pipetted into 0.4 μl cylindrical moulds and allowed to polymerize completely overnight in a humidified chamber to prevent drying out. Prior to experiment, the gels are incubated in PBS for 2 hours to remove loosely bound proteins. Collapse of GVs in the gel utilize the protocol described previously for diffusion characterization.

### Calcium alginate TAPS preparation

Hydrogels were formed using solutions of 2 wt.% alginate (102877-746: VWR) in PBS solutions containing 15 vol.% clustered GVs, 25 mg/mL of etanercept (Y0001969: Sigma-Aldrich) reconstituted in bacteriostatic water as per supplier’s guidelines, calcium carbonate (CaCO_3_) (239216: Sigma-Aldrich) and D-Glucono-δ-lactone (GDL) (8.43794: Sigma-Aldrich). To make a homogenous gel, CaCO_3_ was added to 2 mL of alginate solution to obtain a molarity of 144 mM and vortexed for 10s, and then left to degas at 37 °C overnight. 100 μL of the etanercept solution was then added to the mixture followed by GDL till it doubled the molarity of CaCO_3_ to maintain a neutral pH. The resultant solution is gently mixed to prevent bubbles from being reintroduced and pipetted into 0.4 μl cylindrical moulds and allowed to polymerize completely overnight in a sealed humidified container within an incubator at 25 °C to prevent it from drying out and keep the hydrogels robust. Calcium alginate gels prepared using this method are expected to possess pore sizes of around 5 nm^52^. They are then incubated in PBS for 2 hours to remove loosely bound proteins. The gels were then enterically coated by lightly dip coating them in organic solutions of Eudragit FL30D55 (Evonik Industries) in acetone containing 3 vol.% ethanol and gently blasted with air to dry. The gels were then suspended in PBS prior to administration. This recipe makes enough therapeutic for 2 rats/day at a dose of around 12.5 mg/rat-day. To prepare for oral gavage, a 3 mL syringe was backfilled with 1 mL of PBS. Using a second syringe, TAPS are slowly withdrawn into the gavage needle. The gavage needle is then connected back to the first syringe.

### In vitro release

100 pieces of the acrylamide TAPS were suspended in 30 ml fresh PBS and stirred at 37°C and 100 rpm. The experiment was performed in a dark room to minimize fluorophore photobleaching. Every 2 hours for a total of 12 hours, 10 μL of the suspension was aliquoted into a 90-well plate containing 100 μL of PBS. The wells were then analysed using a Spectramax M5 plate reader with the following wavelength settings: excitation: 587/25 nm and emission: 647/70 nm. After 72 hours, the suspension was aliquoted again to determine the total protein-dye payload present in all ten TAPS cargoes. Fluorescence intensities collected during the time series were then normalised to the final intensity at 72 hours to determine the release kinetics of the TAPS cargoes.

### In vitro spatial targeting

Acrylamide TAPS were evenly distributed and spaced at around 1 mm in a shallow petri dish filled with 1% agarose and allowed to set. An L10-4V 128-element linear array ultrasound transducer (Verasonics) was then operated at peak positive pressure of 200 kPa to image the gels, and subsequently at approximately 600 kPa to trigger the release of fluorophores into the agarose for preselected TAPS.

### In vivo spatial targeting and release

Rats were fasted and given sucrose water, their abdomen hair shaved, and tail cups were placed 17 hours prior to experiments to minimize coprophagy. 1 mL acrylamide TAPS suspension was orally gavaged into the rats. BURST imaging was then performed on the entirety of the rat abdomen 5 hours post-gavage to verify arrival of the GV-gel bolus in the cecum and subsequently trigger the release of fluorophores. BURST images were acquired using a Verasonics Vantage programmable ultrasound system with an L10-4V 128-element linear array ultrasound transducer attached to a custom-made motor stage to scan across the entire abdominal area of the rats. The motor stage was programmed to move the transducer in 0.5 mm steps between transverse planes starting at the bottom of the rib cage and moving towards the tail (approximately 120-160 per rat) and to acquire 2 side-by-side transverse planes to capture the entire width of the abdominal area. To mitigate issues due to tissue motion, images were acquired using a rapid BURST script as described previously^23^ that transmits and acquires three 32-aperature focused beams at a time to improve the frame rate by a factor of 3. The transmit waveform frequency was set to 11.4 MHz to improve the spatial resolution, the focus was set to 12 mm centered on the approximate depth of the cecum, and the number of half-cycles was set to three to increase the BURST signal. BURST images were generated using the temporal-template unmixing algorithm as described previously^23^ on the first three frames which consisted of a pre-collapse frame acquired at 1.6 V and two collapsing frames acquired at 30V. ROIs were manually drawn around the intestines, and BURST acquisitions that occurred during breathing were automatically excluded if the total BURST signal outside the ROI exceeded 162 dB. For display, the BURST signal was summed over the depth of the ROI, excluding pixels below 1 x 10^5^ to reduce background, and the B-mode signal was summed from a depth of 5 to 18 mm. After imaging, the animals were then sacrificed, and the colon harvested and analyzed for fluorescence under a ChemiDoc.

### DSS-Induced ulcerative colitis model

Severe ulcerative colitis was induced in rats allowing rats to consume 4% by weight of dextran sulfate sodium (36-50 kDa colitis-grade DSS, MP Biomedicals, USA) in their drinking water for five days. On day 6, DSS is reduced to 0.4% and TAPS treatment is given once a day for six days until euthanized on day 11. During these 11 days, the rats were monitored daily for changes in weight and stool quality.

### TAPS treatment

During TAPS treatment, the standard laboratory chow is replaced with rodent liquid diet (AIN-76, Bio-Serv) and custom fit tail cups are attached on each rat to prevent coprophagy to improve ultrasound imaging performance. Calcium alginate TAPS loaded with etanercept as previously described were utilized in the TAPS treatment and are freshly prepared prior to daily treatment. It is worth noting that etanercept or human Enbrel is not an FDA approved treatment for colitis, but rather for rheumatoid arthritis^53^. However, we used it as it is a common human anti-TNF-α prescription with significant binding to rodent TNF-α without requiring a surrogate, unlike infliximab^54,55^. 1 mL of the hydrogel suspension was orally gavaged into the rats. 5 hours after administration, the rats are anesthetized and placed on a holder containing an ultrasound transparent window to enable imaging, verify TAPS arrival in the cecum, and trigger GV collapse for etanercept release in the colon using the same imaging parameters described above except a collapsing voltage of 50 was used for 18 frames and a single focused beam was acquired at a time to maximize collapse throughout the GI tract.

### Disease activity index

The DAI was scored using the following criteria: stool consistency (hard: 0, soft: 2, and diarrhea: 4), fecal occult blood using Hemoccult Sensa (Beckman Coulter) (negative: 0, positive: 2, and macroscopic: 4), and decrease in weight relative to the average weight of the healthy controls (less than 1%: 0, 1 to 5%: 1, 5 to 10%: 2, 10 to 20%: 3, and more than 20%: 4).

### Histopathology and scoring

Tissue samples of the colon were fixed in 3.7% paraformaldehyde overnight at 4°C, dehydrated in ethanol and embedded in paraffin. Haematoxylin and eosin (H&E) staining was then performed on the tissues for general histological observation. The histological scores were assessed in a blinded manner for inflammation severity (none: 0, slight: 1, moderate: 2, and severe: 3), polymorphonuclear neutrophil (PMN) infiltration/high power field (HPF) (less than 5: 0, 5 to 20: 1, 21 to 60: 2, 61 to 100: 3, and more than 100: 4), injury depth (none: 0, mucosa: 1, submucosa and mucosa: 2, and transmural: 3), crypt damage (none: 0, basal 1/3: 1, basal 2/3: 2, only surface epithelium intact: 3, and total crypt lost: 4), and adjustment to the tissue involvement multiplied by the percentage factor (0 to 25%: ×1, 26 to 50%: ×2, 51 to 75%: ×3, and 76 to 100%: ×4). The final score was then averaged across all indicators to determine disease severity.

## SUPPLEMENTARY INFORMATION

**Supplementary Table 1.**
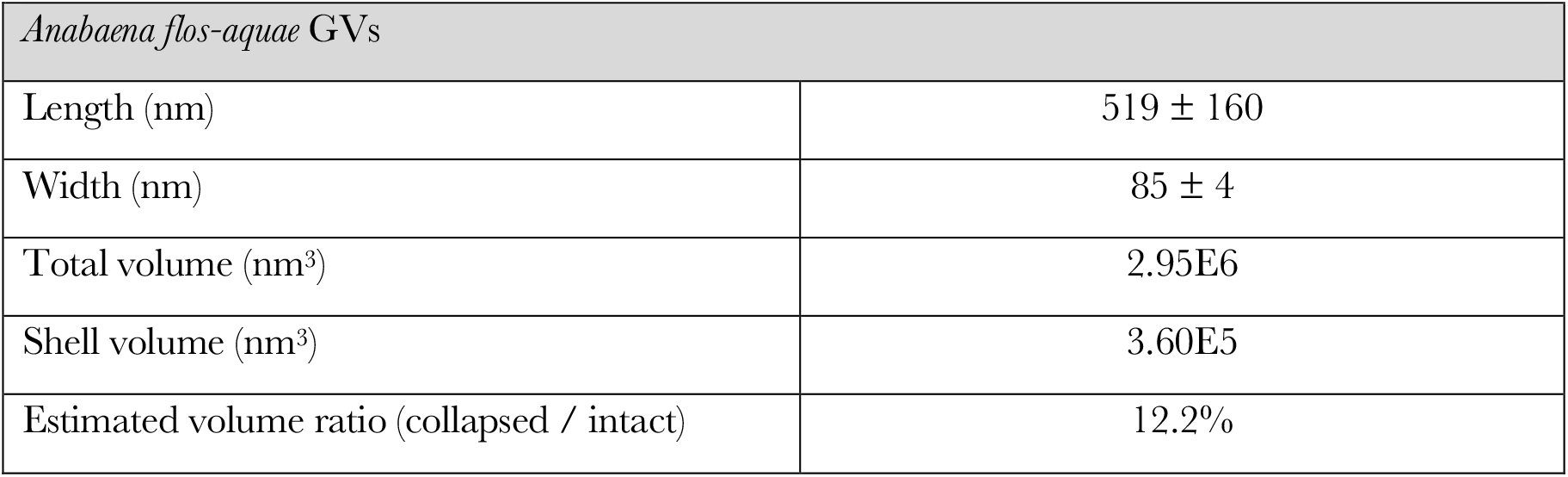
GV dimensions (mean ± s.e.m.) measured using EM for use in volume estimations^13,14^. The GV shell thickness is estimated to be 2.4 nm. Intact GVs are approximated as cylindrical in shape. Collapsed GVs are assumed to be flattened, leaving just the shell.

| *Anabaena flos-aquae* GVs |  |
| --- | --- |
| Length (nm) | 519 ± 160 |
| Width (nm) | 85 ± 4 |
| Total volume (nm<sup>3</sup>) | 2.95E6 |
| Shell volume (nm<sup>3</sup>) | 3.60E5 |
| Estimated volume ratio (collapsed / intact) | 12.2% |

**Supplementary Figure 1.**
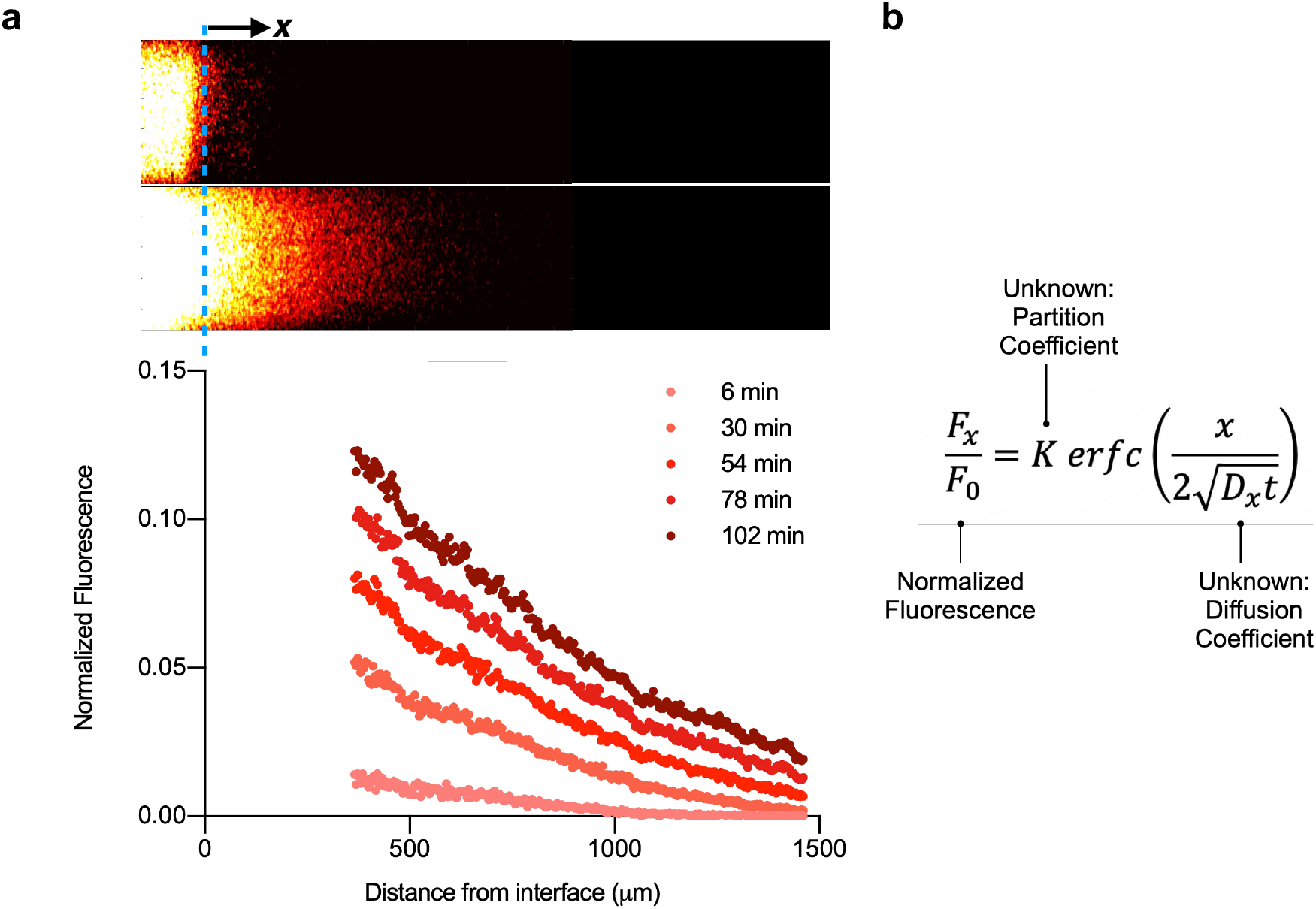
Representative raw data for diffusivity measurements. **a**, Representative fluorescence images at two time points (top) and quantified normalized fluorescence profiles in the capillary diffusion experiment. **b**, Error function used to obtain diffusion coefficients.

**Supplementary Figure 2.**
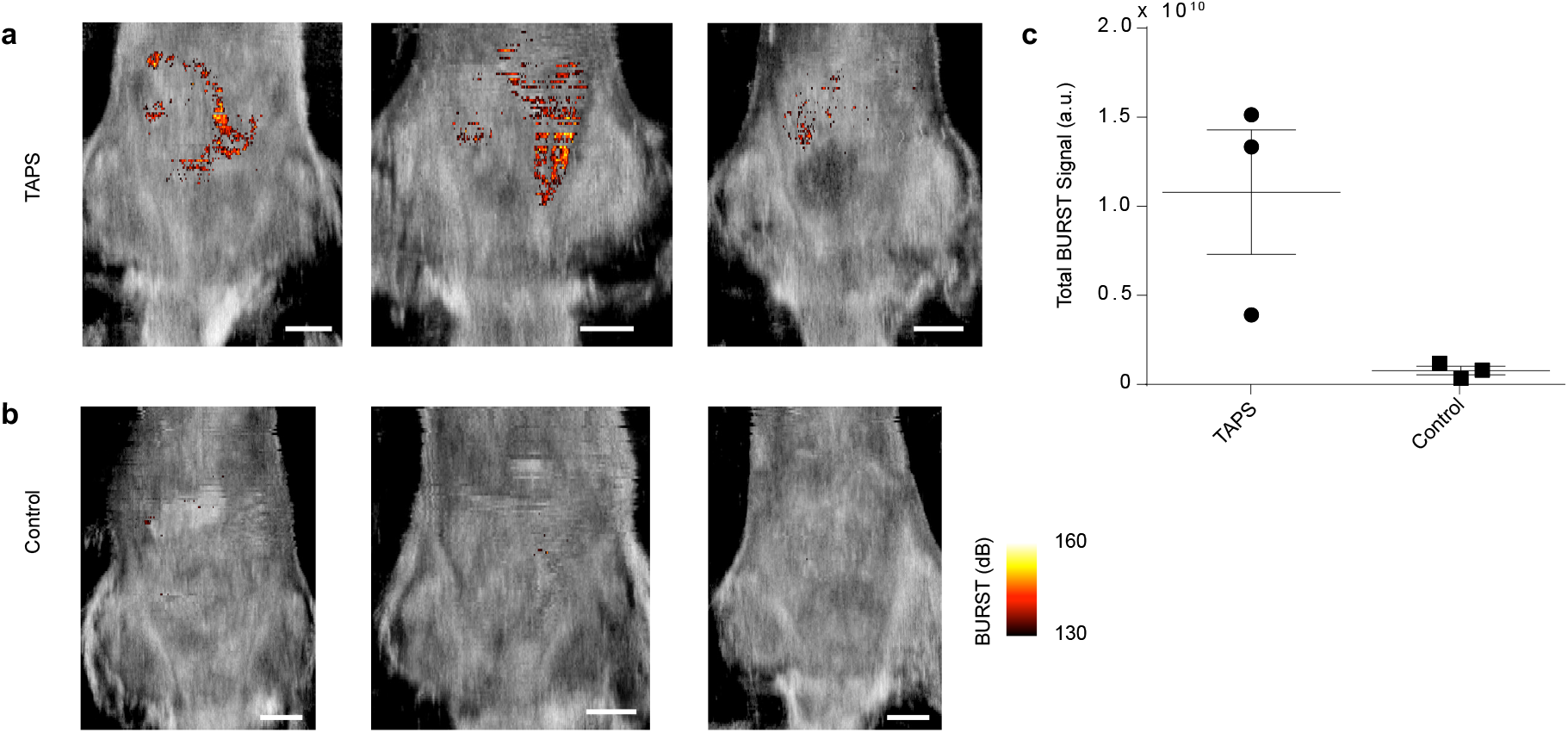
Additional images of TAPS-gavaged and control animals. **a-b**, Additional BURST images of (**a**) rats gavaged with TAPS as described in Fig. 3c and (**b**) controls that did not receive TAPS. **c**, Individual measurements, mean and SEM of total abdominal BURST signal in rats receiving TAPS or controls. Scale bars represent 1 cm.

**Supplementary Figure 3.**
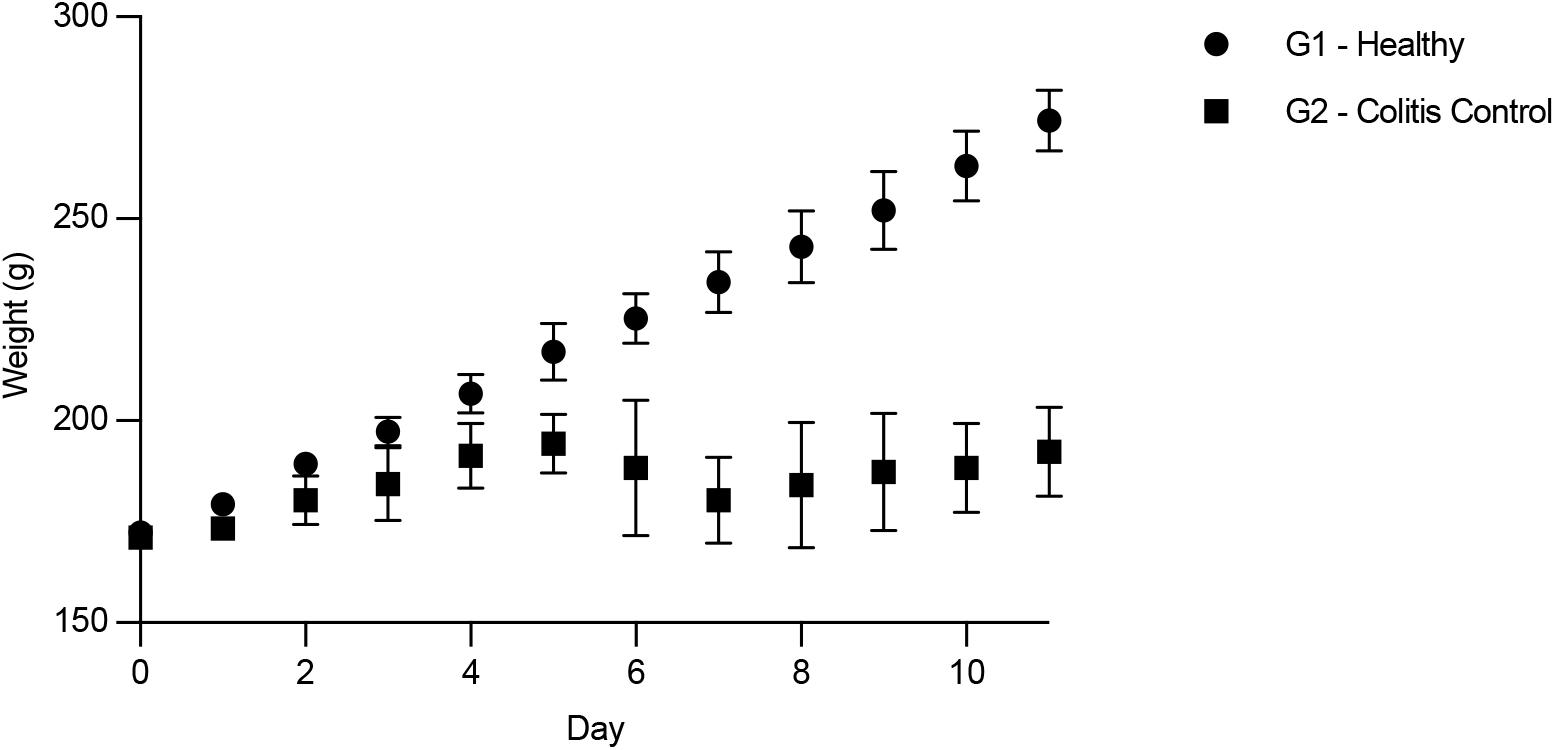
Weight gain trends for healthy and DSS-treated rats. Average weight of rats receiving the DSS treatment described in Fig. 5a or untreated controls.

**Supplementary Figure 4.**
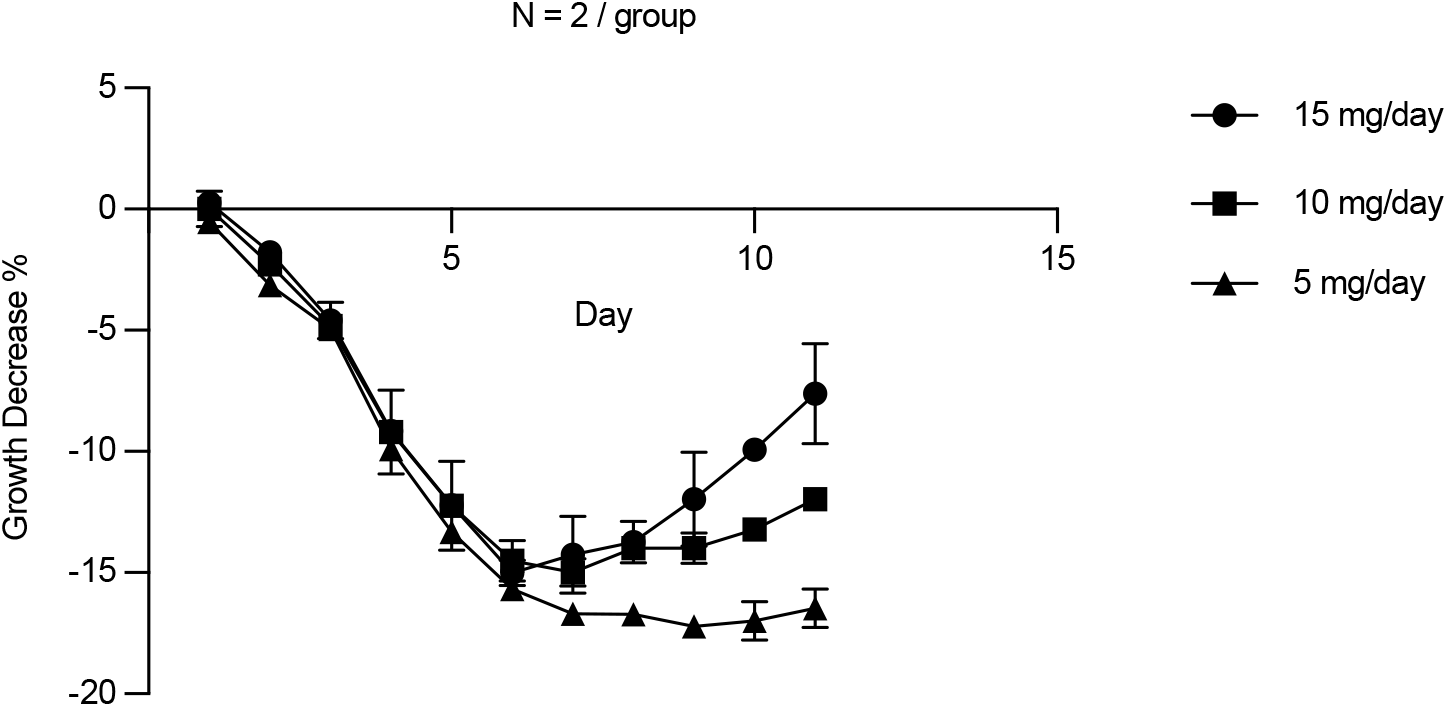
Preliminary dosing experiment for TAPS in colitis model. Weight loss and recovery in rats treated as shown in Fig. 5b with three different doses of TAPS.

